# Efficient memorization of dynamic stimuli with future interactions

**DOI:** 10.1101/2025.09.28.678599

**Authors:** Gonzalo Aparicio-Rodríguez, Daniel Ruiz-Navalón, Paloma Manubens, Abel Sánchez-Jiménez, Carlos Calvo-Tapia, José Antonio Villacorta-Atienza

## Abstract

In nature, survival requires coping with complex time-changing situations in real time. In this process, memory plays a major role since the retrieval of critical information is key to rapid and reliable decision making. This work explores modulation of human memory under the hypothesis that in dynamic scenarios such critical information is encoded as a static map of future interactions. Specifically, the reported results show that dynamic visual stimuli that contain future interactions are better recalled than equivalent stimuli that do not. This is in line with the proposed hypothesis since the former type of stimulus would be encoded in a more simplified way than the latter. Moreover, dynamic stimuli with future interactions are better recalled than simpler dynamic stimuli, which reinforces that the former are processed by a static representation - their map of interactions. This cognitive strategy seems to be modulated by the complexity of the stimulus, since in simple situations differences in recall appear only in men, whereas when complexity increases, such differences do not show gender bias. Therefore, this work proposes an answer to how memory can help us reliably cope with dynamic situations, demonstrating that those critical for survival (such as fighting, chasing, fleeing, etc., which involve interactions) are better remembered, allowing more efficient learning and decision making, essential to deal with our complex and changing world.

---

Memory is an essential cognitive process inextricably linked to spatial encoding of the environment^1^ and, thus, to navigation^2^. In particular, dealing with dynamic situations (which imply the spatial change over time of the subject and/or elements in the environment) requires a central role of memory, as it entails retaining the temporal and spatial location of the elements around the subject^3,4^. In this sense, working memory, besides its relevance in dealing with static scenarios^5,6^, is of paramount importance to overcome the daily challenges of dynamic situations^7^, playing a key role in maintaining situation awareness (i.e. the ability to perceive and interpret a situation correctly to respond accordingly)^8^, allowing visual integration and representation of mobile object characteristics in humans^9^, and providing a more efficient and safer navigation in dynamic environments when implemented to artificial intelligences^10^.

A critical aspect of working memory is its restricted capacity, as only a limited amount of information can be retained for a limited time^11^. Thus, aspects such as energy, time and structural efficiency are crucial for memory^12,13^ and consequently information complexity often needs to be reduced for a proper encoding and recalling^14,15^. This is especially important in time-changing situations, which pose specific challenges to working memory^9^, since dynamic stimuli comprise complex information, as the subject needs to process his own and the other elements’ position, directions, velocities and accelerations for each moment. Therefore, in dynamic situations complexity reduction is mandatory to encode and retrieve information in real time for an efficient decision making^16^.

The information complexity in the real world imposes constraints not only on memory, but on processing in general. In living beings, processing their immediate environment is one of the basic capabilities required for survival^17^, which is particularly relevant in dynamic scenarios, where opportunities and threats often come from other individuals. To explain how humans are capable to cope with time-changing environments in a fast and reliable way, it has been proposed that our brain optimizes the processed information by internally representing the dynamic situation as a static map of the possible future interactions between the involved elements (encounters, collisions, etc.), including the subject itself^18^. This process, called time compaction, does not therefore consider other spatiotemporal attributes of the dynamic situation, such as initial positions, velocities, trajectories of the moving elements, etc., so it avoids the explicit representation of the temporal dimension by embedding time into space.

Recent studies have shown that human cognition makes use of time compaction to process dynamic stimuli^19^. Behavioral experiments have shown that rats can encode dynamic visual stimuli by mapping predicted interactions between the perceived moving elements^20^. Furthermore, bats’ hippocampus encodes future positions of themselves in potential collision points^21^. However, although it has been shown that time compaction is part of the cognitive strategies involved in processing and understanding the world, the role of this mechanism on - and whether it influences-other cognitive functions such as learning^16^ and memory remains unclear.

The case of memory is particularly relevant, since memorization and recall are fundamental requirements in survival-critical tasks, such as navigation^22^. Furthermore, information reduction provided by other processes observed in memory operate similarly to information reduction in time compaction^23^. This similarity suggests exploiting the relationship between stimulus simplicity and recall efficiency^24,25^ to explore the role of this cognitive mechanism in human working memory. Following time compaction hypothesis, a dynamic stimulus where future interactions exist will be internally represented by spatially arranging such predicted interactions in a static map, called compact internal representation or CIR^18^ (Fig. 1A). On the other hand, a stimulus with no future interactions cannot be compacted into a CIR and will be encoded through the diverse spatiotemporal attributes of the moving elements (initial positions, directions, trajectories, velocities, etc.; Fig. 1B). Thus, since the stimulus with future interactions will be encoded in a simplified way through its CIR, it will be memorized using less cognitive resources than a similar dynamic stimulus with no interactions. Consequently, it is to be expected that, once memorized, the former are more stable than the latter^24,25^.

**Figure 1.**
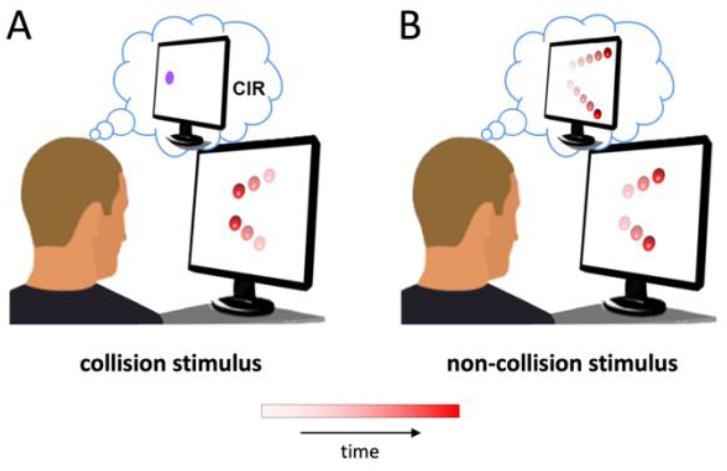
Experimental hypothesis following time compaction. **A**. When a stimulus consisting of two moving red balls moving towards the same point (collision stimulus) is shown to a participant, it is compacted as a CIR, thus, only the zone where the collision would take place (purple dot) is represented. Note that the balls disappear before collision, so such collision is not displayed. **B**. When a stimulus in which two moving red balls diverge (non-collision stimulus) is displayed, the stimulus is not internally compacted and is encoded as a spatiotemporal event. The movement of the balls is depicted using a color scale ranging from light red to red, where lightest indicates the first position of the balls and red indicates their last position, as shown in the legend bar (the same for the following figures).

The results obtained here are aligned with this theoretical prediction, revealing that in a recognition memory task, dynamic stimuli with future interactions are better recalled than those without interactions. Interestingly, this effect has only been observed in men during simple tasks, but in both women and men during complex tasks, which is consistent with previously reported gender differences in the salience of time compaction^19^. These results support the significance of time compaction as a novel mechanism in human cognition.

## Experimental design

To study the role of time compaction in working memory and test the above mentioned hypothesis, we employed a behavioral methodology based on the classic recognition memory task^26,27^. The visual stimuli considered consisted of two moving red balls, and were of two types: those in which the balls move to collide, with no collision displayed, and those in which the moving balls do not collide. They are collision and non-collision stimuli, respectively.

In the first phase of the test, called encoding phase, participants were shown a set of collision and non-collision stimuli, which will be called old stimuli, and they were asked to memorize them (Fig. 2A). After this encoding phase, a short video unrelated to the experiment was shown to participants before starting the next stage, called recalling phase. During the recalling phase, participants were shown a set of old and new (not previously displayed) stimuli. In this stage, after each stimulus, participants were asked if they had already seen it, answering by clicking with the mouse on a yes or no button on the screen, and their answers were recorded (Fig 2B).

**Figure 2.**
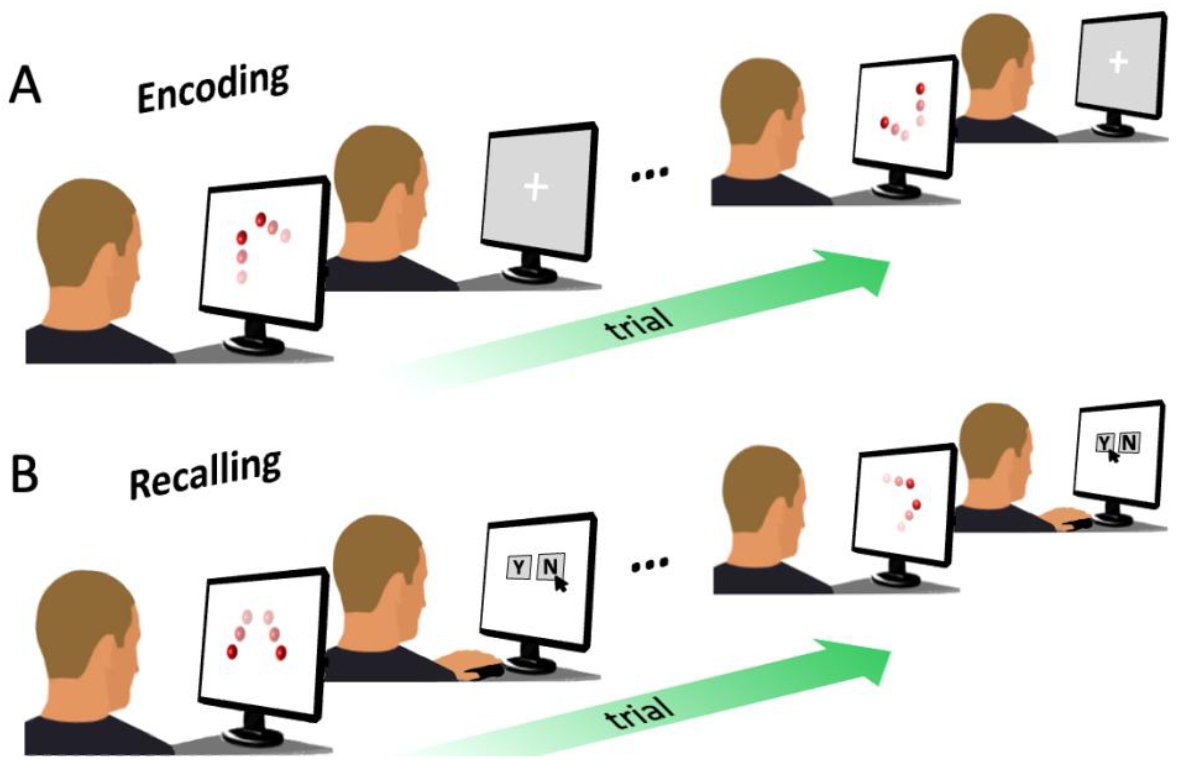
Structure of the experimental procedure, consisting of two consecutive phases. **A**. In the encoding phase, participants were asked to memorize a set of stimuli, which will be called old stimuli. Each stimulus display was followed by a grey screen with a static white Greek cross on its center, so participants could discriminate between the different stimuli. **B**. In the recalling phase, a set of old stimuli and a set of new stimuli were displayed. After each stimulus, the participant was asked if they had already seen that stimulus in the encoding phase.

Based on this design, we tested two experimental conditions of increasing complexity on two participant samples. These experimental conditions were called simple and complex recognition memory tests. We also conducted another recognition memory test, called looming recognition memory test, in whose stimuli only one moving ball appeared and, thus, did not include future interactions. This test was performed to evaluate the hypothesis against basic dynamic stimuli (a single moving ball) and to discard possible additional effects from sources other than time compaction, which could have biased the results of the other tests.

### Simple recognition memory test

In the simple recognition memory test, the balls in the stimuli always moved in an apparent bidimensional plane. In the old stimuli, the trajectories of the red balls were such that their sum vector coincided with the bisector of one of the quadrants of the xy plane (Fig. 3A). On the contrary, in the new stimuli the sum of the vector trajectories of the moving balls resulted in a vector parallel to the x or y axis (Fig. 3B). Thus, the simple recognition memory test was designed as an easy task, since new stimuli were clearly distinguishable from old stimuli.

**Figure 3.**
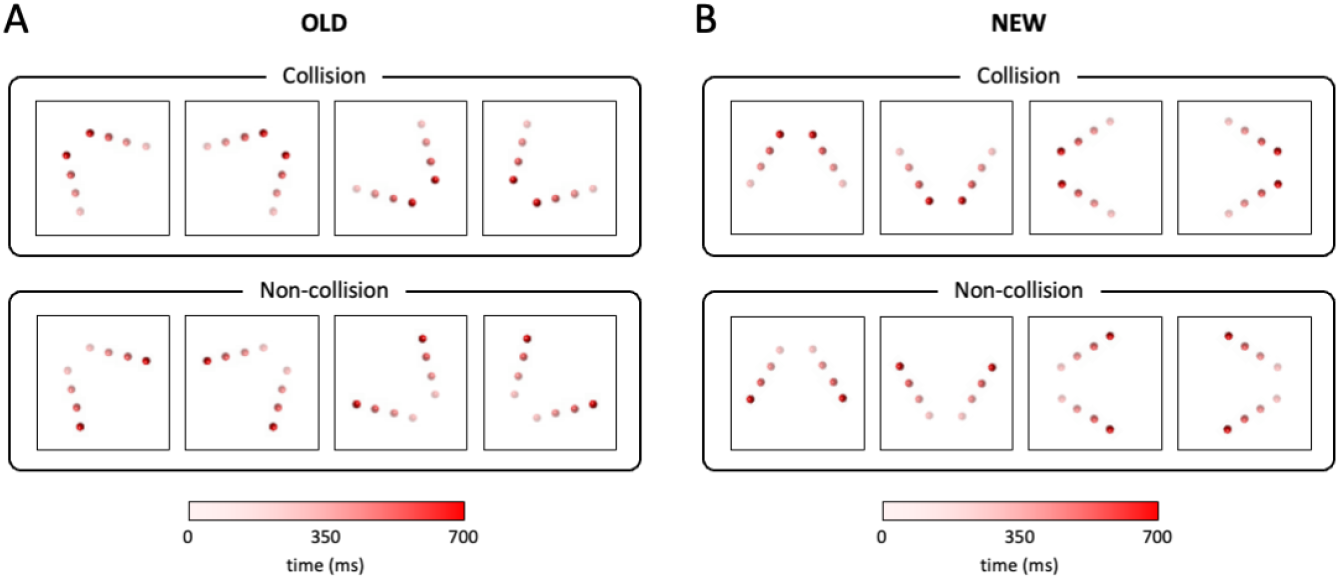
Dynamic stimuli shown in the simple recognition memory test. **A**. Old stimuli. The sum of motion vectors of the balls results in vectors bisecting the four quadrants of the xy plane. Upper row depicts the four collision stimuli and bottom row depicts the four non-collision stimuli. **B**. New stimuli. The vectorial sum of the trajectories of the balls results in vectors parallel to the x or y axis. Upper and lower rows depict the four collision and four non-collision stimuli, respectively.

During the encoding phase, the complete set of 8 possible stimuli were shown in a pseudorandom order (4 collision and 4 non-collision; Fig. 3A). During the recalling phase, a random set of 3 familiar collision, 3 familiar non-collision (Fig. 3A), 3 novel collision and 3 novel non-collision stimuli (Fig. 3B) was displayed with a random order of appearance.

### Complex recognition memory test

In the complex recognition memory test, stimuli consisted of the two moving red balls, but now their movement took place in a 3-dimensional space, i.e. red balls trajectories also varied along the z axis. This way, the balls in the collision/non-collision stimuli moved away from/towards the participant, so they got smaller/bigger. Additionally, in the complex recognition memory test differences among stimuli are more subtle than in the simple recognition memory test (i.e., ball’s directions vary more gradually among stimuli), so they are intended to be more difficult to distinguish (see Methods). Contrary to the simple recognition memory test, in which the old and new stimuli were distinguishable, in the complex recognition test there is no differential key to distinguish between both types of stimuli. This trait is intended to make this test more complex for participants to elicit the use of the CIR versus other cognitive strategies, as supported by time compaction hypothesis.

During the encoding phase, a random set of 4 collision and 4 non-collision stimuli was shown in random order to each participant. In the recalling phase, the complete set of 16 possible stimuli was shown to participants, meaning that 8 collision stimuli (4 of them old stimuli and 4 of them new stimuli) and 8 non-collision stimuli (4 of them old stimuli and 4 of them new stimuli) were displayed (Fig. 4). The order of appearance of the stimuli in the recalling phase was randomized.

**Figure 4.**
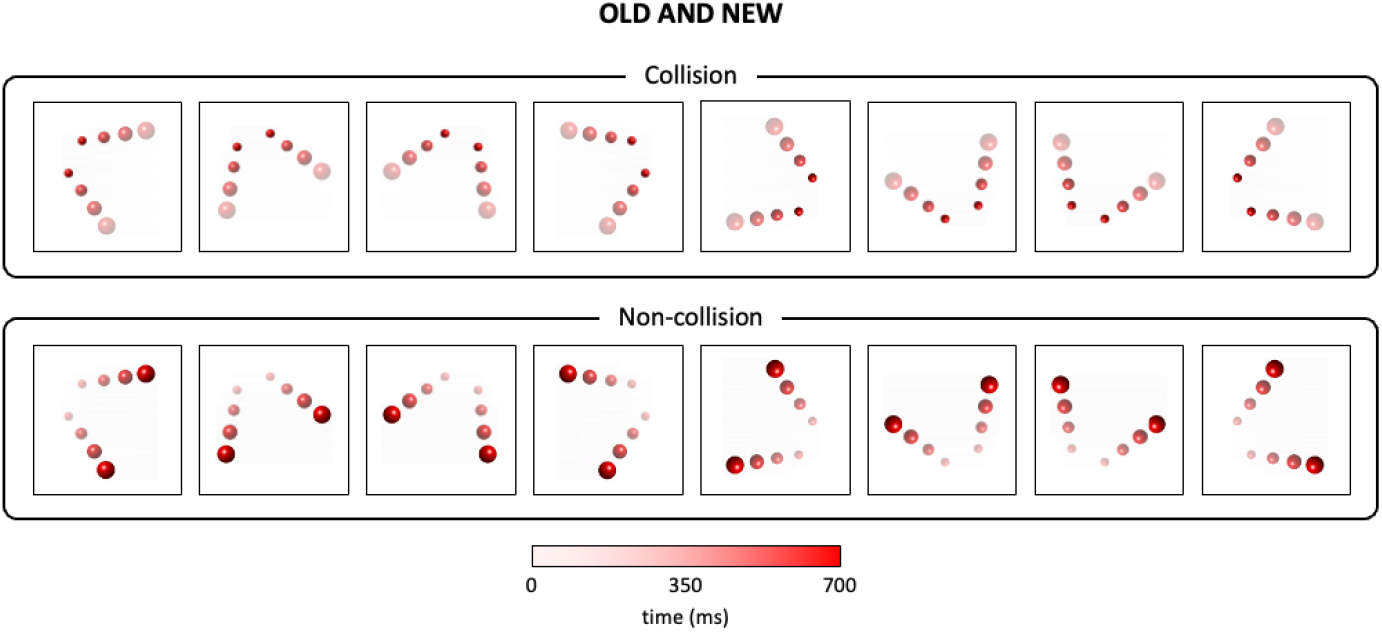
Dynamic stimuli shown in the complex recognition memory test. As in the simple recognition memory test, collision and non-collision stimuli were displayed. The complexity is intended to make it more difficult to distinguish the different stimuli, and it is introduced by 1) including the third dimension by changing the size of the balls as they move, 2) gradually changing the orientation of the trajectories of the red balls, avoiding horizontal and vertical orientations, and 3) randomly drawing old and new stimuli from the same set of collision and non-collision stimuli.

### Looming recognition memory test

The working hypothesis implies that a collision stimulus, being represented similarly to a static stimulus through its CIR, should be better remembered than a simpler dynamic stimulus. To explore this prediction, we performed the looming recognition memory test, which is the same as the complex recognition memory test but replacing the stimuli in Fig. 4 by equivalent simpler stimuli with one moving ball (Fig. 5). Furthermore, this test allows us to check whether the looming effect can be ruled out as a cognitive factor contributing to the potential differences between collision and non-collision stimuli, since approaching stimuli capture the subject’s attention^28^.

**Figure 5.**
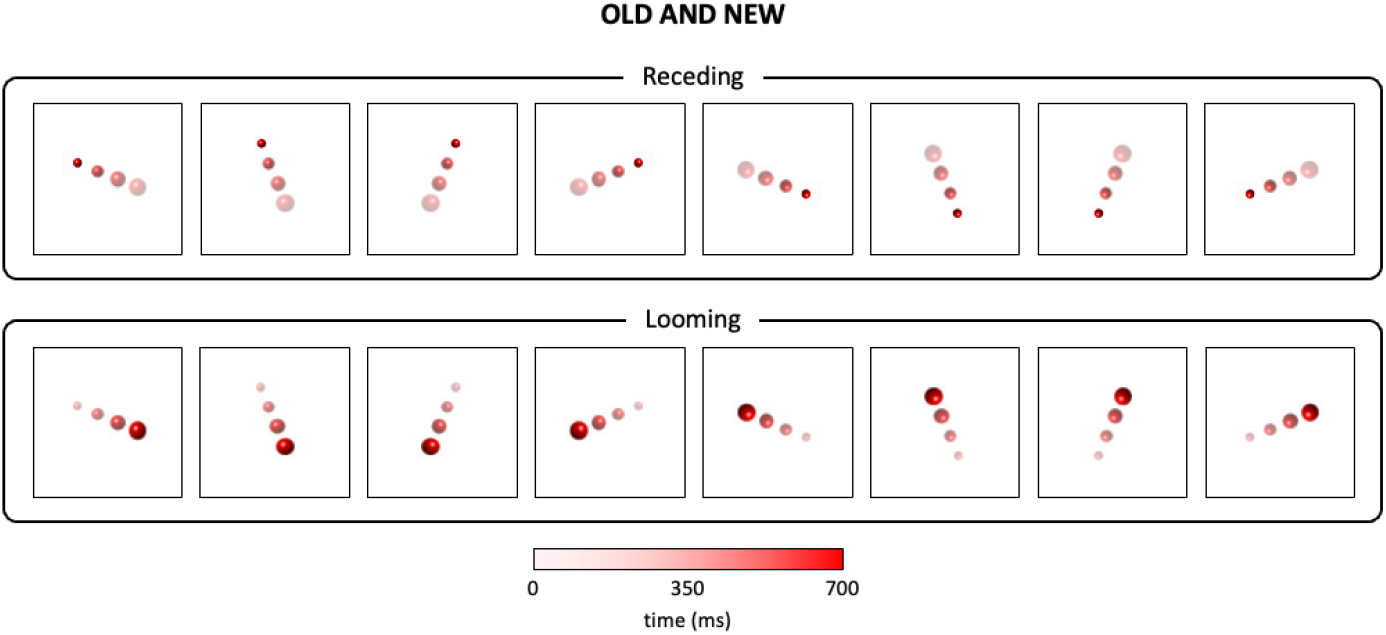
Dynamic stimuli shown in the looming recognition memory test. They have the same structure as the stimuli for the complex recognition memory test, except only one red ball is displayed, so the two types of stimuli are called receding and looming.

The looming recognition memory test stimuli, shown in Fig. 5, consisted of only one moving red ball instead of two, so no compaction of the situation was possible. In these stimuli, the ball also moved in a 3-dimensional space, and two different types of situations were considered: looming stimuli, in which the ball moved towards the participant (i.e. ball got bigger), and receding stimuli, in which the ball moved away from the participant (i.e. ball got smaller). Note that, with this configuration, each stimulus in the complex recognition memory test has its correspondence in one of the looming recognition memory test stimuli. For example, the first collision stimulus of the complex recognition memory test shown in Fig. 4 corresponds to the first stimulus of the looming recognition memory test in Fig. 5 in terms of orientation, position, ball behavior, etc. Regarding the structure of the test (trajectory orientations; absence of differential key between old and new stimuli; number of stimuli displayed during encoding and recalling phases; randomization of the stimulus sequences; etc.), it was identical to the complex recognition memory test.

## Results

### Simple recognition memory test: Men recall collision stimuli better than non-collision

The working hypothesis, stating that collision stimuli are better recalled than non-collision stimuli, was initially tested in the simple recognition memory test. On the one hand, regarding the effect of having previously seen a stimulus in the encoding phase, participants obtained better results in the recalling phase when considered new stimuli rather than when considered old stimuli (GLMM p-value = 0.005, Table SXXX5; GLMM Odd Ratio (OR) = 0.63, Table S6). As mentioned in the Materials and Methods section, new stimuli were designed to be easy to differentiate from the old stimuli due to their orientation and lack of profundity. Therefore, the new stimuli served as a baseline to check that participants performed the simple recognition memory test correctly in terms of attention and motivation. Also, no differences between genders or stimuli types were observed when only new stimuli were analyzed (GLMM p-value = 0.126 and p-value = 0.351, respectively; Table S7).

On the other hand, regarding old stimuli, men and women got significantly different results on the test (GLMM p-value = 0.003; Table S8). While men got greater success when collisions were shown (GLMM p-value = 0.001 and Odd Ratio collisions vs non collisions in men (OR) = 3.05; Table S9), women had similar performances no matter a collision or non-collision stimulus was displayed (p-value = 0.744; Table S10) (Fig. 6). The differences found were not affected by the experimental conditions (p-value = 0.171; Table S11).

**Figure 6.**
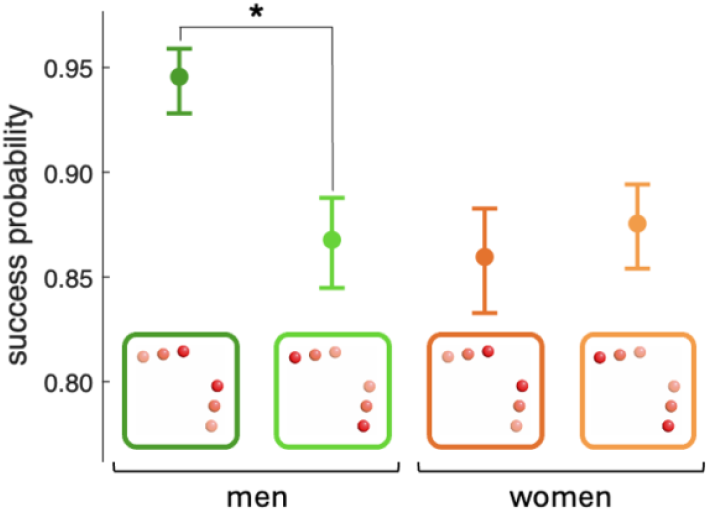
Performance in the recalling phase of the simple recognition memory test for old stimuli by gender and stimulus type. Points and error bars show punctual and interval estimations (standard errors), respectively, obtained from the probabilities of the computed models for each gender and stimulus type. Schematics for stimulus type are depicted in the color boxes below, with the red balls moving in convergent and divergent trajectories to represent collision and non-collision stimuli, respectively (the movement of the balls is represented according to the color scale in the figures above). Thus, dark and light green colors stand for collision and non-collision stimuli shown to men, respectively, and dark and light orange stands for collision and non-collision stimuli displayed to women, respectively. Asterisks show significant differences between groups.

No effect of trial on success was found (GLMM p-value = 0.364; Table S12). Thus, as might be expected from the simplicity of the task, no tiredness effect and no learning or forgetting process took place throughout the test. No effect of other stimulus features (e.g. orientation of the trajectories of the balls) on success was either found besides being collision or non-collision stimulus (GLMM p-value = 0.108; Table S13).

### Complex recognition memory test: Both men and women remember collision stimuli better than non-collision stimuli

The gender differences reported above are in line with the results that demonstrated the existence of time compaction in humans^19^. We conjecture that these differences appear in simple tasks, as they can be addressed by resorting to different strategies. However, we hypothesize that in complex dynamic situations the advantages of time compaction may push both women and men to resort to this cognitive strategy, making the gender differences in the simple recognition memory test disappear. This hypothesis is explored in the complex recognition memory test.

Visualization of the stimuli during the encoding phase (i.e. old or new stimulus) is a significant factor in explaining successful recall, as old stimuli yield better results than new stimuli (GLM p-value = 0.031; Table S14; OR = 1.85; Table S15). These results differ from the simple recognition memory test, in line with the increased complexity of the task. Unlike the simple recognition memory test, in the complex recognition memory test the new stimuli have no structural differences with old stimuli and make it more difficult for participants to decide whether they have already seen them in the encoding phase or not.

Regarding the old stimuli, success probability in recall significantly depended again on the type of stimulus, but no differences were found between men and women (GLM p-value = 0.320; Table S16). In the complex recognition memory test, the better recall of collision stimuli versus non-collision ones was modulated by the trial in which the stimulus appeared (GLM p-value = 0.036; Table S17). In the first analyzed trial collision stimuli are better recalled than non-collision stimuli (Fig. 7A; GLM p-value = 0.002; Table S17), with a 38% higher probability of success (random answer as baseline; OR = 2.57). Then, success probability for collision stimuli progressively decreases with trial (Fig. 7B; GLM p-value = 0.002; Table S18), showing the expected forgetting process. On the other hand, no such effect of trial is observed in non-collision stimuli (GLM p-value = 0.853; Table S19). Note that by the end of the recalling phase both types of stimuli are equally remembered. Again, no effect of other stimulus features (besides being a collision or a non-collision stimulus) on success was found (GLMM p-value = 0.241; Table S20).

**Figure 7.**
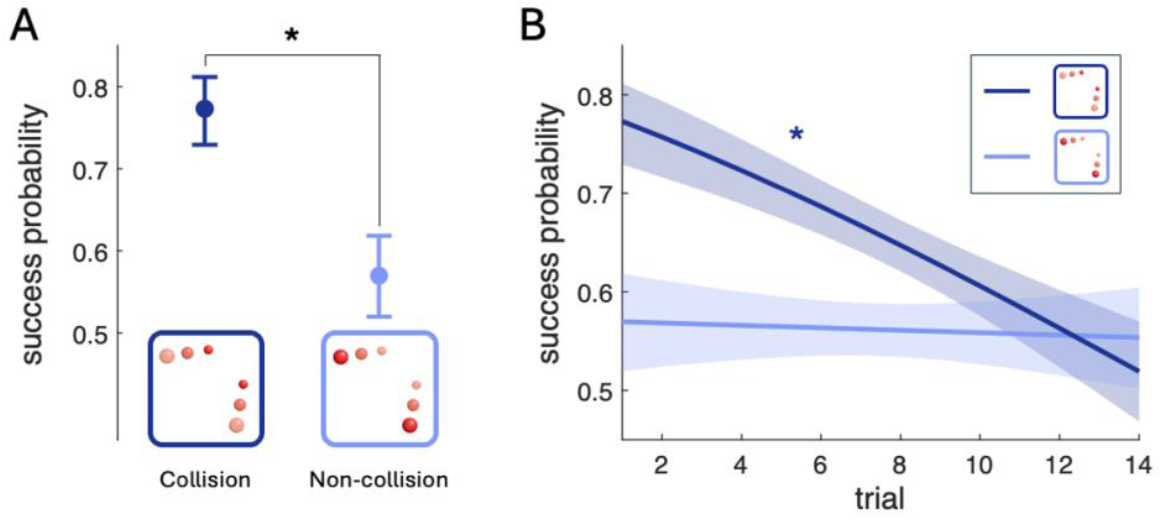
Performance during the recalling phase of the complex recognition memory test for old stimuli. **A**. Performance by stimulus type in the first trial. Points show punctual estimations and error bars show interval estimations (standard errors). Schematics for stimulus type are displayed in the color boxes below, representing the stimulus type (collision and non-collision as in Fig. 4). Asterisk shows significant differences. **B**. Temporal performance by stimulus type. Lines represent the generalized linear model for each type and shadowed areas the standard error for each of the models (dark and light blue stand for collision and non-collision stimuli, respectively). The asterisk denotes a significant trial dependence.

### Looming recognition memory test: Better recall of complex collision stimuli than of simpler dynamic stimuli

The static representation of dynamic stimuli proposed by time compaction suggests that the dynamic collision stimuli themselves should be better remembered than simpler dynamic stimuli. This conjecture is the objective of the looming recognition memory test.

As in the previous experiment, participants obtained better results when they were shown an old stimulus rather than a new stimulus (GLM p-value = 0.001; OR = 1.43; Table S21). Among the old stimuli, there is no significant effect of stimulus type (i.e. being a looming or a receding stimulus) nor participant gender (Fig. 8A) (GLM p-value = 0.938 and p-value = 0.795, respectively; Table S22). Thus, the absence of differences in the looming recognition memory test suggests that the differential effect between collision and non-collision stimuli in the complex recognition memory test is not related to the looming effect.

**Figure 8.**
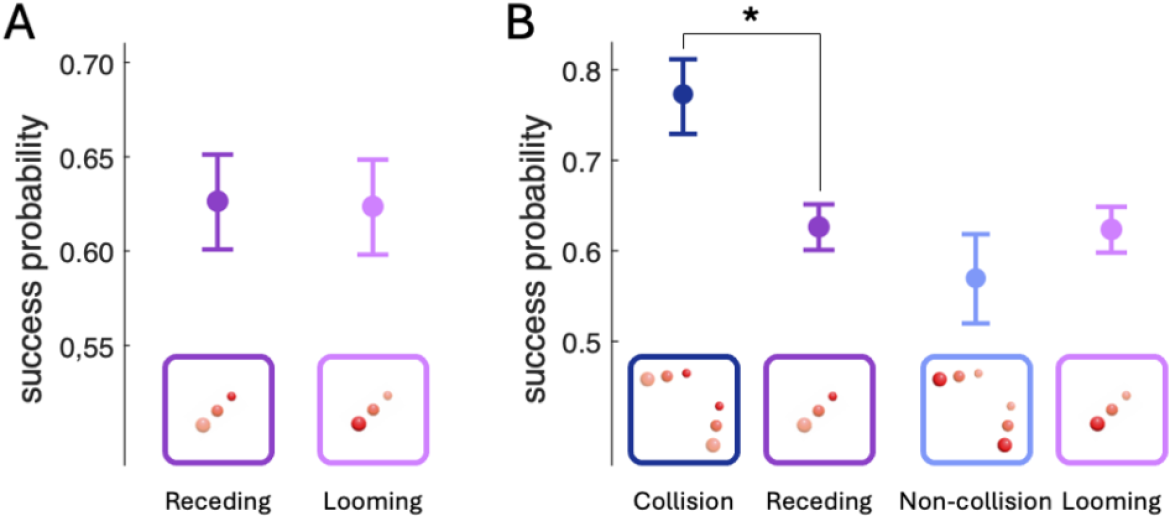
Performance in the looming recognition memory test and comparison with complex recognition memory test. **A**. Recalling phase performance in the looming recognition memory test for old stimuli by stimulus type. **B**. Comparison of success probability during recalling in the first trial between complex (Fig. 7A) and looming recognition memory tests (panel A). Asterisk denotes significant differences for the relevant comparison: collision vs. receding stimuli (other comparisons not shown). Points and error bars show punctual estimations and standard errors, respectively. Schematics for stimulus type are displayed in the color boxes below.

On the other hand, the equivalence between stimuli in complex and looming recognition memory tests (see Materials and Methods section) allows going deep into the hypothesis of time compaction. Complex and looming stimuli only differ in their level of complexity, as the number of moving objects is different. Therefore, it would be expected that the recall of looming receding stimuli (dark purple in Fig. 8A) would be better than that of collision complex stimuli (dark blue in Figs. 7), since the former are simpler (one vs. two moving balls). However, for the first trial (before the effect of forgetting shown in Fig. 7B) the recall of the collision stimuli in the complex recognition memory test is significantly better than that of the corresponding stimuli (receding) in the looming recognition memory test (GLM p-value = 0.018, Table S23) (dark blue and purple in Fig. 8B). Hence, when a dynamic stimulus contains future interactions, it is better recalled than a simpler dynamic stimulus. Finally, the orientations of the trajectories of the ball on each stimulus had no effect on success (GLM p-value = 0.781; Table S24).

## Discussion

Dynamic situations are a primordial part of animals’ natural environment and memorizing and recalling them properly is crucial for survival. The internal representation of a time-changing situation by a spatial map of the predicted impending interactions (the CIR), proposed by time compaction hypothesis, suggests a mechanism to optimally encode, learn and memorize dynamic environments^16^. In particular, the role of memory is especially important from a survival perspective, since in rapidly changing threat situations (fight, hunt, flee, etc.), fast and reliable access to critical information is mandatory for real-time decision making.

The results of the simple recognition memory test show that when a stimulus contains future interactions and thus, according to the hypothesis, can be internally represented in a compacted way (the CIR), it is better recalled than when the stimulus does not contain impending interactions. Furthermore, the results obtained in the complex recognition memory test for collision stimuli are not only better than those obtained for the non-collision stimuli from the same test, but also better than those obtained for the looming recognition memory test. The stimuli in this test contain only a moving element instead of two, comprising less information.

Therefore, it would be expected that these stimuli would be easier to memorize and recall than the stimuli in the complex recognition memory test. This paradoxical result can be explained from the perspective of time compaction. According to this hypothesis, a collision stimulus is encoded as a static representation mapping the future collision point, i.e., its CIR (Fig. 1A). Thus, the CIR, being static, is simpler than the corresponding dynamic stimulus in the looming recognition memory test. For this reason, it is to be expected that the probability of recalling the CIR, and thus the corresponding complex dynamic stimulus, is higher than that of recalling the looming recognition memory test stimuli, as revealed by the experiment.

### Memory and time compaction

The compaction of the temporal information into the spatial dimension provides an optimization of the processing of those scenarios in which interactions could exist, focusing on the relevant information, i.e., when and where encounters could take place. On the other hand, those scenarios with no future interactions, being less relevant to the subject, do not need to be compacted. This dichotomy between different dynamic situations provides a testable approach to the role of time compaction in working memory: dynamic visual stimuli with impending interactions, being encoded in a simpler way, will require less cognitive resources and will therefore be more stable during memorization and recalling than dynamic stimuli with no future interactions. Our results confirm this working hypothesis, showing a better recall of stimuli that can be compacted than those that cannot.

This suggests that time compaction would allow storing experienced dynamic situations in a way that makes them easy to be retrieved and manipulated to react in real time and behave flexibly to the current situation^29^. This process would occur in many situations of daily life, both for humans (e.g. walking through a crowded street) and animals (e.g. hunting a prey or escaping from a predator), and thus it would be essential for survival. Furthermore, our results show that this optimization of memorization based on time compaction also occurs when the subject does not intervene in the situation, thus allowing observational learning. This kind of learning is critical for survival, as humans and animals can benefit from experiences without endangering themshelves^30,31^.

What mechanisms underly the differences in memorization and retrieval of compacted and non-compacted situations? We conjecture that the observed results root at the engram level. Engrams are physical traces of memory in the shape of neuronal circuits with potentiated synaptic connections^32–35^ that are hypothesized to contain the internal representations of memory^36^. Indeed, the traditional cognitive map is thought to rely on engrams for its memorization and retrieval^37,38^, as place cells function as physical traces of memory^22,39^ (although the exact role of different place cells subpopulations is yet unclear^39–41^). We hypothesize that differences between compacted and non-compacted stimuli must be related to engram codification. Allocation of engrams (i.e. which neurons take part of the neuronal ensemble) is determined by the excitability of the neuron candidates, as more excitable neurons out-compete their less excitable neighbors and then comprise the engram^42^. This excitability modulates a continuous process of reshaping the engram as a process of learning/forgetting^43^. Following time compaction hypothesis, since a dynamic stimulus with future interactions is encoded by the spatial mapping of these interactions, the neural ensemble recruited for the engram associated with this compact representation will be smaller and more specific than that linked to non-compacted stimuli, as they require the encoding of multiple spatiotemporal aspects, as initial positions, trajectories, velocities, etc. Therefore, the first type of memories would be less susceptible to the interference from other stimuli, as the competition to allocate the engram would be less tough, resulting in greater memory stability during memorization and forgetting, in line with the results reported.

This differential efficiency in the memorization of dynamic situations with and without predicted interactions could also have implications for the consolidation of memories. There are different mechanisms to construct long-term memories^44^, as consolidation of the working memory through similar experiences^45^ and through replay^46,47^. Therefore, time compaction could be an additional mechanism to be considered in the formation of long-term memory, facilitating the consolidation of experiences in dynamic situations where interactions are a priority and thus helping to navigate and perform actions in time-changing environments^29^.

The essence of time compaction is the embedding of the temporal information into the spatial dimension (the ‘when’ is transformed into the ‘where’^18^). The connection between this transformation and memory here explored is relevant because of the close bond between memory and spatial encoding, widely described in the literature. For instance, time cells in the hippocampus, which provide a temporal framework for spatial events and encode additional spatial aspects of the environment too, are thought to provide a structural order to episodic memory^48^. Another example is hippocampal replay, which is involved in the processing of past experiences but also plays a role in spatial encoding and navigation^49^. Some authors argue that memory is a salience of spatial cognition^50^, while others argue that spatial cognition is a collateral effect of memory^51^. Despite this, there is consensus that the cortical region where both processes take place is the hippocampal formation^37,52,53^, an area also well known for allocating engrams together with other cortical regions, mainly the medial prefrontal cortex^39^. In this sense, the role of the hippocampal formation as the region underpinning time compaction has not yet been explored, but it has been hypothesized to be critically involved in this mechanism^20^. In fact, experiments in bats suggest that hippocampal place cells are implicated in time compaction^21^ and it has been shown that time compaction arises from simple network conditions agnostic to any cortical regions but coherent with hippocampal circuits^54^. Another plausible candidate neurons mediating time compaction in the hippocampal formation are the object vector cells, a population of cells located in the medial entorhinal cortex that fire at specific distances and orientations from objects^55^.

Based on these theoretical and experimental results, we suggest a central role of hippocampus in the processing of dynamic situations based on the mapping of future interactions. It has been hypothesized that such a map of predicted interactions generalizes to dynamic situations the concept of cognitive map, usually related to static environments^56^. Spatial cognition is based on the encoding of essential characteristics of the environment, such as the position of the subject in space^22^, the size or metric of the environment^57^, the location of borders^58^ and objects^55^, etc., The coordinated activity of the responsible cell populations leads to the cognitive map, an internal representation of the environment supporting memory and decision making during navigation^22,59,60^. The information reduction provided by the cognitive map by encoding relevant information in static scenarios is challenged in dynamic environments, as the information in temporal dimension could dramatically increase the processing load. The mapping of future interactions along with other significant spatial attributes allows eliminating time and encoding the dynamic scenario as a static representation, the CIR, generalizing the concept of cognitive map. This could contribute to explain how animals and humans are able to make real-time decisions in rapidly changing environments, which is mandatory for survival^16^.

### Gender differences in cognitive strategies

This work shows differential results in working memory performance in terms of gender differences. In the simple task, men outperformed women in recalling old collision stimuli, whereas both genders were equally successful in recalling old non-collision stimuli. This is aligned with results obtained in previous study, which showed that while there is saliency of time compaction in men, women seem to resort to a broader set of cognitive strategies^19^. However, when the task complexity is increased, such as in the complex recognition memory test, no gender differences were found. These gender biased observations are not uncommon regarding working memory tests. Gender differences in visuospatial working memory tests are frequently reported^61^ and even different brain circuits appear to be involved depending on the gender of the participant^62^. However, although men more often outperform women on visuospatial working memory tasks^63–65^, the gender being favored by these eventual differences depends on multiple factors, including the type of task to be performed^61,66^. Various authors hypothesize that these differences rely on the cognitive strategy used to solve the task^19,67,68^. This explanation aligns with our results, since different individuals can use different approaches to solve a task^69,70^ and a single individual may change their strategy depending on performance^71^ or context^72^.

Consistent with these results, the gender dependence of recall performance and the cognitive strategies used during the experiments reported here can be interpreted in terms of task complexity. When the faced task is simple, men are more prone to resort to time compaction as encoding strategy to adapt to dynamic situations, whereas women rely in a broader set of strategies^19^. However, when the task complexity increases, both women and men equally rely on time compaction to deal with difficult time-changing situations. This is compatible with the main idea of time compaction hypothesis: optimizing the processing of complex dynamic scenarios to make real-time decisions by engaging critical cognitive mechanisms, such as learning and memory.

Nevertheless, the reasons underlying the different preferences in cognitive strategies have not yet been unraveled, since experiments on the action of sexual hormones in working memory point to unclear results^73,74^ and the effect of socio-cultural factors cannot be discarded. It is also noteworthy that other possible explanations for the gender differences in working memory have also been proposed, such as performing tests with computers, with which male participants seem to feel more comfortable^75^, better spatial skills in male participants^76^ and negative stereotypes around women’s performance in spatial tasks^67^. Therefore, future work should also address which factors and mechanisms underlie the gender differences in time compaction recurrence.

This paper shows that the memorization of dynamic visual stimuli in which there are imminent and predictable interactions are more stable, and therefore better recalled, than those without future interactions, and even more stable than simpler dynamic stimuli. This result is compatible with the predictions of time compaction hypothesis on the memorization of dynamic situations, since recognition memory test have proven to be a suitable proxy for the study of memory^77^. Specifically, these findings contribute to explain how, after adequately learning to cope with rapidly changing situations (through training, playing, observation, etc.), humans are able to make real-time decisions to deal with these familiar scenarios (or even similar ones), since the corresponding CIRs can be accurately retrieved from memory and converted into actions, as they contain the appropriate information to interact with the environment^16^. The fact that women, more versatile regarding the use of cognitive strategies in dynamic situations^19^, resort to time compaction in complex contexts suggests that this may be a relevant strategy in human cognition. Furthermore, recent results reported in bats^21^ and rats^20^ imply that these processing mechanisms involving encoding, learning, memory and decision making in dynamic environments could extend to other animal species. This suggests that time compaction could be a cognitive invariant in different animal species, playing an important role among brain processes critical for survival of living beings.

## Methods

### Experiment technical details

The experimental design was implemented in PC using R software^78^ and its extension Shiny^79^, through the RStudio environment^80^, which allowed us to perform the tests telematically to a great number of participants. To validate this data collection and rule out any bias coming from this, we also performed the simple recognition memory test in person using the software MATLAB^81^. Both software received the same specifications for the implementation.

In both encoding and recalling phases, the presentation of each stimulus consisted of two consecutive repetitions of the ball movements, with a duration of 1 s per repetition. During the first 0.7 s the two moving red balls were displayed, disappearing for the remaining 0.3 s. The presentation of a stimulus was followed by a grey screen with a white Greek cross in the center displayed for 3 s. This latter screen served as a separator between stimuli and forced the participant to center the view in the middle point of the screen, so all stimuli were visualized starting from the same point. Shadows and shines were added to the balls to give them realism and ensure that participants would perceive their future interactions as collisions and not as two plain circles crossing each other. In the simple recognition memory test, to prevent biases when recalling collision or non-collision stimuli, we randomly showed each participant one of two pseudorandom sequences of old stimuli that alternated between collision and non-collision stimuli (see Supplementary). In the complex and looming recognition memory tests, the order of appearance of the stimuli was completely randomized.

In the complex and looming recognition memory test, regarding the motion in the xy plane, the vectorial sum of the trajectories of the balls in the complex recognition memory test and the ball’s trajectory in the case of the looming recognition memory test, resulted in vectors oriented 22.5°, 67.5°, 112.5°, 157.5° 202.5°, 247.5°, 292.5 and 337.5° with respect to the x axis.

The answer buttons from the recalling phase appeared at the center of the screen next to each other and their left or right position was randomized between participants. Once the recalling phase was completed, participants were requested to introduce their gender and age, which were the only personal information collected.

At the beginning of the experiment, participants were shown a text on the computer screen with the information needed to perform the memory test, avoiding providing any other information that could bias the results (e.g., no mention of potential collisions). This information was also read aloud by the researcher. Participants were given only the following instructions: 1) the experiment was a memory test and it comprised two parts, 2) in the first phase, a set of stimuli consisting in two moving balls will be displayed and should be memorized, and 3) in the second phase, they were going to see stimuli and after each one they will be asked if they have already seen them. Any questions from participants were answered by referring to these instructions.

### Participants

The simple recognition memory test was performed by 237 participants who completed 2370 trials (1180 observations/trials from 66 women and 52 men, for the R software setup; and 1190 observations/trials from 70 women and 49 men, for the MATLAB software setups). For the complex recognition memory test, 99 volunteers participated in the experiment completing 1386 trials (51 women and 48 men). Finally, the looming recognition memory test was performed by 104 participants who completed 1456 trials (67 women and 37 men). Participant age ranged from 18 years old to 41 years old, with similar distributions for both genders and for each test and setup (Tables S1, S2, S3 and S4).

All participants for each test were different (no person participated in more than one test) and they provided informed consent to anonymously take part in the experiments, according to the experimental procedures approved by the Institutional Review Board (Committee of Bioethics, National Distance Education University). All experiments were performed following the guidelines and regulations set forth by the Declaration of Helsinki.

### Statistical analysis

We analyzed simple, complex and looming recognition memory tests separately. First two trials of each test were intended to serve as an adaptation to the setup and task, and were not included in the analysis. We performed analysis of success (answering correctly during the recalling phase) by fitting individual probabilities of successful answers to random intercept mixed models with logit link (i.e. logistic regression) (GLMMs), using the individual (i.e. participant) as the grouping variable (i.e. upper level). When the variance related to the upper level was not large enough to justify the use of mixed models, we used generalized linear models (GLMs) instead. The initial set of variables analyzed were gender, stimulus type (i.e. collision or non-collision stimulus for the simple and complex recognition memory tests; looming and receding stimulus for the looming recognition memory test) and whether it was an old or new stimulus. From a model with these variables and their interactions, we followed a backwards stepwise elimination procedure to select the set of variables with explanatory power following p-value criterion. When interactions between variables were found, we disentangled them by analyzing and interpreting the interaction terms separately. Models fit was performed via maximum likelihood. We used Bobyqa controller^82^ when needed, to avoid convergence problems.

We also analyzed the influence of trial (i.e. ordinal position in which the stimulus is shown), checking for significant interactions of trial and stimulus with gender, stimulus type and whether the stimulus is an old or a new stimulus. In order to separately estimate the effect of these factors on each trial we used pairwise comparisons to the selected model. The effect of stimulus (i.e. each individual stimulus, with its particular orientation of ball trajectories) on success was also assessed through the same procedure used for the effect of trial, to check that there was no bias coming from this factor.

For every model, we checked residuals and distributions’ assumptions. For all mixed models used, intraclass correlation indexes (ICCs) were large enough to justify the use of multilevel regression (see Supplementary). All statistical analyses were performed by means of R software under RStudio IDE^78^ using the libraries lme4^83^ (computation of mixed models) and emmeans^84^ (pairwise comparisons of the selected model).

## Acknowledgments

This research was supported by the Ministry of Science and Innovation (Spain), Grant PID2022-138659NB-I00 and the European Social Fund Plus (ESF+), Grant PEJ-2024-AI/SAL-GL-32302.

