## Supplementary material for "Efficient memorization of dynamic stimuli with future interactions"

### Simple recognition memory test

The encoding phase of the simple recognition test consisted in the display of one of two pseudorandom sequences of stimuli to be memorized. These two sequences are as follow: for the first sequence, non-collision with both balls moving from the right bottom corner, collision with both balls moving towards the top right corner, non-collision with both balls moving from the top left corner, collision with both balls moving towards the bottom left corner, non-collision with both balls coming from the upper right corner, collision with both balls moving towards bottom right corner, non-collision with both balls moving from the bottom left corner and collision with both balls moving towards the upper right corner; for the second sequence collision with both balls moving towards the upper right corner, non-collision with both balls moving from the right bottom corner, collision with both balls moving towards the top right corner, non-collision with both balls moving from the top left corner, collision with both balls moving towards the bottom left corner, non-collision with both balls coming from the upper right corner, collision with both balls moving towards bottom right corner and non-collision with both balls moving from the bottom left corner.

### Tables with statistical results

Glossary of variables included in statistical analysis

The following have been included in the models below:

- phase: dichotomic variable indicating the phase of first appearance of the stimulus: Old for those stimuli appearing for the first time in the encoding phase and new for those stimuli appearing in the recalling phase.
- type: dichotomic variable indicating whether the stimulus is a collision or a non-collision stimulus for the recognition memory test and recognition memory test or whether it is a receding or looming stimulus for the looming recognition memory test.
- gender: dichotomic variable indicating the declared gender of the participant: men or women.
- trial: variable indicating the order of appearance of the stimulus during the recalling phase.
- success: dichotomic variable indicating whether the participant provided a correct or a wrong answer: 1 for correct answers and 0 for wrong answers.

**Table S1:** Frequencies and age description of participants of simple recognition memory test in the R setup by gender.

| Gender | Absolute frequency | Relative frequency | Mean age | Median age | Quantile 0.05 for age | Quantile 0.95 for age |
|---|---|---|---|---|---|---|
| Men | 52 | 0.44 | 21.12 | 19 | 18 | 29 |
| Women | 66 | 0.56 | 18.79 | 18 | 18 | 27.25 |

**Table S2:** Frequencies and age description of participants of simple recognition memory test in the MATLAB setup by gender.

| Gender | Absolute frequency | Relative frequency | Mean age | Median age | Quantile 0.05 for age | Quantile 0.95 for age |
|---|---|---|---|---|---|---|
| Men | 49 | 0.41 | 21.02 | 19 | 18 | 29.6 |
| Women | 70 | 0.59 | 20.8 | 19 | 18 | 31 |

**Table S3:** Frequencies and age description of participants of complex recognition memory test by gender.

| Gender | Absolute frequency | Relative frequency | Mean age | Median age | Quantile 0.05 for age | Quantile 0.95 for age |
|---|---|---|---|---|---|---|
| Men | 48 | 0.485 | 23.08 | 22 | 18 | 35.95 |
| Women | 51 | 0.515 | 20.82 | 21 | 18 | 28.5 |

**Table S4:** Frequencies and age description of participants of looming recognition memory test by gender.

| Gender | Absolute frequency | Relative frequency | Mean age | Median age | Quantile 0.05 for age | Quantile 0.95 for age |
|---|---|---|---|---|---|---|
| Men | 12 | 0.324 | 22.67 | 23 | 18.275 | 30.18 |
| Women | 25 | 0.676 | 21.8 | 22 | 19 | 29.4 |

The following tables contain the main information about the mixed models and GLMs used for the interpretations and graphs of the results section. Columns in the tables describe from left to right the predictor variable or covariate, the covariate estimation, its odd ratio (calculated by $e^{\text{estimate}}$), odd ratio 95% confidence interval, and the p-value signification. For binary variables the covariate column indicates in between brackets the comparison category, in contrast to the non-appearing category, which would be the reference category. For the reference category, covariate = 0, while for the comparison category, covariate = 1. This means that estimate is only applied to the comparison category (the one in between brackets) (e.g. if the reference category of gender is "women" and the odd ratio = 2, this means that the ratio for women is twice the ratio for men). Those p-values lower to 0.05 (and thus, considered significant) appear in bold format. For mixed models, tables also include information below about intraclass correlation coefficient or ICC, variance of the upper level ($\tau_{00}$), number of individuals used as upper levels, and number of observations. For GLMs, tables also include information below about the number of observations.
As the variable stimulus (particular stimulus shown, *i.e.* orientation of the trajectories of the ball) involves 8 categories for the simple recognition memory test and 16 categories for the complex and looming recognition memory test (as there are 8 and 16 different old stimuli for these tests, respectively), we checked the significance of the factor from the GLMM/GLM using the ANOVA test from the R library car[85]. The below tables regarding this factor show the output from this test.

For the simple recognition memory test, every participant answered correctly at least 4 out of the 10 trials analyzed and a total of 224 participants (94.51%) succeeded in between 7 and 10 trials. This proves the accessibility of the test and the easiness to identify the old stimuli.

Regarding the complex recognition memory test, participants answered correctly to between 3 and 11 trials, out of a total of 14 analyzed trials. This worse distribution of results compared to the simple recognition memory test shows the increased difficulty of the complex experiment. In this sense, perhaps due to such complexity, there was no inter-individual effect on success in the task.

Finally, for the looming recognition memory test, participants correctly answered to between 3 and 12 trials, out of a total of 14 analyzed trials. Again, there was no inter-individual effect in succeeding in the test.

**Table S5:** simple recognition memory test: success GLMM including all variables.

| Covariate | Estimate | Odd ratio (OR) | OR Confidence Interval* | Sig. |
|---|---|---|---|---|
| (Intercept) | 2.81 | 16.56 | 9.54 – 28.76 | **<0.001** |
| gender [Women] | -0.66 | 0.52 | 0.27 – 0.99 | **0.046** |
| type [Collision] | -0.09 | 0.91 | 0.46 – 1.79 | 0.786 |
| phase [Old] | -1.02 | 0.36 | 0.20 – 0.66 | **0.001** |
| gender [Women] × type [Collision] | 0.45 | 1.57 | 0.68 – 3.64 | 0.293 |
| gender [Women] × phase [Old] | 0.82 | 2.28 | 1.07 – 4.84 | **0.032** |
| type Collision] × phase [Old] | 1.16 | 3.18 | 1.29 – 7.83 | **0.012** |
| (gender [Women] × type [Collision]) × phase [Old] | -1.60 | 0.20 | 0.07 – 0.62 | **0.005** |

$\tau_{00}$ id = 0.69 (n = 237); ICC = 0.17; 2370 observations

*α = 0.05

**Table S6:** simple recognition memory test: success GLMM for phase.

| Covariate | Estimate | Odd ratio (OR) | OR Confidence Interval* | Sig. |
|---|---|---|---|---|
| (Intercept) | 2.48 | 11.94 | 9.26 – 15.39 | **<0.001** |
| phase [Old] | -0.47 | 0.63 | 0.48 – 0.82 | **<0.001** |

$\tau_{00}$ id = 0.65 (n = 237); ICC = 0.16; 2370 observations

*α = 0.05

**Table S7:** simple recognition memory test: success GLMM for new stimuli.

| Covariate | Estimate | Odd ratio (OR) | OR Confidence Interval* | Sig. |
|---|---|---|---|---|
| (Intercept) | 3.00 | 20.00 | 10.81 – 37.02 | **<0.001** |
| type [Collision] | 0.20 | 1.22 | 0.80 – 1.85 | 0.351 |
| gender [Women] | -0.45 | 0.64 | 0.36 – 1.14 | 0.126 |

$\tau_{00}$ id = 1.64 (n = 237); ICC = 0.33; 1158 observations

*α = 0.05

**Table S8:** simple recognition memory test: success GLMM for old stimuli.

| Covariate | Estimate | Odd ratio (OR) | OR Confidence Interval* | Sig. |
|---|---|---|---|---|
| (Intercept) | 1.79 | 5.98 | 4.00 – 8.93 | **<0.001** |
| type [Collision] | 1.03 | 2.79 | 1.54 – 5.07 | **0.001** |
| gender [Women] | 0.18 | 1.19 | 0.72 – 1.97 | 0.489 |
| type [Collision] × gender [Women] | -1.10 | 0.33 | 0.16 – 0.69 | **0.003** |

$\tau_{00}$ id = 0.70 (n = 237); ICC = 0.18; 1212 observations

*α = 0.05

**Table S9:** simple recognition memory test: success GLMM for old collision and men.

| Covariate | Estimate | Odd ratio (OR) | OR Confidence Interval* | Sig. |
|---|---|---|---|---|
| (Intercept) | 2.05 | 7.73 | 4.44 – 13.47 | **<0.001** |
| type [Collision] | 1.11 | 3.05 | 1.62 – 5.71 | **0.001** |

$\tau_{00}$ id = 1.69 (n = 101); ICC = 0.34; 517 observations

*α = 0.05

**Table S10:** simple recognition memory test: success GLMM for old stimuli and women.

| Covariate | Estimate | Odd ratio (OR) | OR Confidence Interval* | Sig. |
|---|---|---|---|---|
| (Intercept) | 1.84 | 6.29 | 4.45 – 8.90 | **<0.001** |
| type [Collision] | -0.07 | 0.93 | 0.61 – 1.42 | 0.744 |

$\tau_{00}$ id = 0.30 (n = 136); ICC = 0.08; 695 observations

*α = 0.05

**Table S11:** simple recognition memory test: success GLMM including all variables - non-significant effect of platform.

| Covariate | Estimate | Odd ratio (OR) | OR Confidence Interval* | Sig. |
|---|---|---|---|---|
| (Intercept) | 2.94 | 18.88 | 10.49 – 33.98 | **<0.001** |
| setup [R] | -0.24 | 0.79 | 0.56 – 1.11 | 0.171 |
| gender [Women] | -0.67 | 0.51 | 0.27 – 0.98 | **0.043** |
| type [Collision] | -0.10 | 0.90 | 0.46 – 1.77 | 0.763 |
| phase [Old] | -1.03 | 0.36 | 0.20 – 0.65 | **0.001** |
| gender [Women] × type [Collision] | 0.45 | 1.57 | 0.68 – 3.64 | 0.292 |
| gender [Women] × phase [Old] | 0.82 | 2.28 | 1.07 – 4.83 | **0.032** |
| type Collision] × phase [Old] | 1.17 | 3.22 | 1.31 – 7.95 | **0.011** |
| (gender [Women] × type [Collision]) × phase [Old] | -1.60 | 0.20 | 0.07 – 0.62 | **0.005** |

$\tau_{00}$ id = 0.67 (n = 237); ICC = 0.17; 2370 observations

*α = 0.05

**Table S12:** simple recognition memory test: success GLMM for new stimuli - non- significant effect of trial.

| Covariate | Estimate | Odd ratio (OR) | OR Confidence Interval* | Sig. |
|---|---|---|---|---|
| (Intercept) | 2.23 | 9.26 | 5.53 – 15.50 | **<0.001** |
| trial | -0.03 | 0.97 | 0.92 – 1.03 | 0.364 |

$\tau_{00}$ id = 0.69 (n = 237); ICC = 0.17; 1212 observations

*α = 0.05

**Table S13:** simple recognition memory test: ANOVA test for success GLMM model used to analyze the factor stimulus among the old stimuli.

| Covariate | Chisq. | Df | Sig. |
|---|---|---|---|
| stimulus | 11.769 | 7 | 0.108 |

**Table S14:** complex recognition memory test: selected success glm.

| Covariate | Estimate | Odd ratio (OR) | OR Confidence Interval* | Sig. |
|---|---|---|---|---|
| (Intercept) | -0.13 | 0.88 | 0.71 – 1.09 | 0.230 |
| type [Collision] | 0.38 | 1.46 | 1.08 – 1.98 | **0.013** |
| phase [Old] | -0.10 | 0.90 | 0.67 – 1.21 | 0.493 |
| type [Collision] × phase [Old] | 0.47 | 1.60 | 1.04 – 2.46 | **0.031** |

1386 observations

*α = 0.05

**Table S15:** complex recognition memory test: selected success glm in function of phase.

| Covariate | Estimate | Odd ratio (OR) | OR Confidence Interval* | Sig. |
|---|---|---|---|---|
| (Intercept) | -0.18 | 0.83 | 0.72 – 0.96 | **0.015** |
| phase [Old] | 0.61 | 1.85 | 1.49 – 2.29 | **<0.001** |

1386 observations
*α = 0.05

**Table S16:** complex recognition memory test: success glm with trial-type interaction for old stimuli - non-significant effect of gender.

| Covariate | Estimate | Odd ratio (OR) | OR Confidence Interval* | Sig. |
|---|---|---|---|---|
| (Intercept) | 0.36 | 1.43 | 0.94 – 2.19 | 0.100 |
| type [Collision] | 0.95 | 2.59 | 1.41 – 4.81 | **0.002** |
| trial | <-0.01 | 1.00 | 0.94 – 1.05 | 0.889 |
| gender [Women] | -0.16 | 0.85 | 0.63 – 1.16 | 0.320 |
| type [Collision] × trial | -0.08 | 0.92 | 0.85 – 0.99 | **0.033** |

692 observations
*α = 0.05

**Table S17:** complex recognition memory test: success glm with trial-type interaction for old stimuli.

| Covariate | Estimate | Odd ratio (OR) | OR Confidence Interval* | Sig. |
|---|---|---|---|---|
| (Intercept) | 0.28 | 1.32 | 0.89 – 1.98 | 0.166 |
| type [Collision] | 0.94 | 2.57 | 1.40 – 4.77 | **0.002** |
| trial | -0.01 | 1.00 | 0.94 – 1.05 | 0.853 |
| type [Collision] × trial | -0.08 | 0.92 | 0.85 – 0.99 | **0.036** |

692 observations
*α = 0.05

**Table S18:** complex recognition memory test: selected success glm for old collision stimuli.

| Covariate | Estimate | Odd ratio (OR) | OR Confidence Interval* | Sig. |
|---|---|---|---|---|
| (Intercept) | 1.23 | 3.41 | 2.17 – 5.49 | **<0.001** |
| trial | -0.09 | 0.92 | 0.86 – 0.97 | **0.002** |

345 observations
*α = 0.05

**Table S19:** complex recognition memory test: selected success glm for old non-collision stimuli.

| Covariate | Estimate | Odd ratio (OR) | OR Confidence Interval* | Sig. |
|---|---|---|---|---|
| (Intercept) | 0.28 | 1.32 | 0.89 – 1.98 | 0.166 |
| trial | -0.01 | 1.00 | 0.94 – 1.05 | 0.853 |

347 observations
*α = 0.05

**Table S20:** complex recognition memory test: ANOVA test for success GLM model used to analyze the factor stimulus among the old stimuli.

| Covariate | Chisq. | Df | Sig. |
|---|---|---|---|
| stimulus | 18.433 | 15 | 0.241 |

**Table S21:** looming recognition memory test: selected success glm.

| Covariate | Estimate | Odd ratio (OR) | OR Confidence Interval* | Sig. |
|---|---|---|---|---|
| (Intercept) | 0.15 | 1.17 | 1.01 – 1.35 | **0.038** |
| phase [Old] | 0.36 | 1.43 | 1.16 – 1.76 | **0.001** |

1456 observations
*α = 0.05

**Table S22:** looming recognition memory test: selected success glm for old stimuli.

| Covariate | Estimate | Odd ratio (OR) | OR Confidence Interval* | Sig. |
|---|---|---|---|---|
| (Intercept) | 0.53 | 1.70 | 1.27 – 2.28 | **<0.001** |
| gender [Women] | -0.04 | 0.96 | 0.70 – 1.31 | 0.795 |
| type [Receding] | 0.01 | 1.01 | 0.75 – 1.37 | 0.938 |

728 observations

*α = 0.05

**Table S23:** complex recognition memory test vs looming recognition memory test: effect of the experiment for collision/receding stimuli on the first trial.

| Covariate | Estimate | Odd ratio (OR) | OR Confidence Interval* | Sig. |
|---|---|---|---|---|
| (Intercept) | 1.95 | 7.00 | 2.75 – 23.64 | **<0.001** |
| experiment [Looming] | -1.54 | 0.21 | 0.05 – 0.72 | **0.018** |

62 observations

*α = 0.05

**Table S24:** looming recognition memory test: ANOVA test for success GLM model used to analyze the factor stimulus among the old stimuli.

| Covariate | Chisq. | Df | Sig. |
|---|---|---|---|
| stimulus | 10.585 | 15 | 0.781 |

<u>Participants' response time</u>

For the simple recognition memory test, participant response times (i.e. time taken since the stimulus disappeared till the moment when the yes or no bottom was clicked) were recorded in the MATLAB setup.

The analysis of response time was performed using linear mixed models (with lme4 package[83]; P-values for lineal mixed models were obtain using lmertest[86]). As for the logistic mixed models mentioned in the main text, first two trials from the recalling phase were excluded. Analysis was performed by fitting individual response times to random intercept mixed models, using individual as the upper level. As response times did not follow a normal distribution, to assess signification of variables we transformed response times observations using a Box-Cox transformation[87], so model assumptions were not violated. Effect size assessment was performed by using the untransformed variables. The initial set of variables analyzed were success, gender, stimulus type and whether the stimulus was an old or a new stimulus. From a model with these variables and their interactions, we followed a backwards stepwise elimination procedure to select the set of variables with explanatory power. When interaction between variables were found, we disentangled them by analyzing and interpreting the interaction terms separately in function of one of the other interaction term's group.

We also analyzed the influence of trial and stimulus in response time, to check that there was no bias coming from these factors. We did this by fitting again success rates and Box-Cox transformed response times to random intercept mixed models, using individual as the upper level. We also checked for significant interactions of trial and stimulus with gender, stimulus type and whether the stimulus was an old or a new stimulus.

All residuals and distributions' assumptions were checked. For all mixed models used, intraclass correlation indexes (ICCs) were large enough to justify the use of multilevel regression.

From the above-mentioned factors, the only one having a significant influence in participants' response time was success (p-value <0.001; Table S25). Specifically, correct answers were given 564 ms earlier than wrong answers (Table S26). While no effect of stimulus (besides being a collision or a non-collision stimulus) on response time was found, trial did have a significant effect on response time (Table S25), rendering for each consecutive trial a 32 ms faster response than in the prior trial (Table S26).

The following tables contain the main information about the mixed models selected in the response time analysis. Reference category is equally indicated as in the models above. For the transformed mixed models, tables also include information below about intraclass correlation coefficient or ICC, variance of the upper level ($\tau_{00}$), variance of residuals ($\sigma^2$), number of individuals used as upper levels, and number of observations. For GLMs, tables also include information below about the number of observations.

**Table S25:** Selected transformed response time mixed model.

| Covariate | Estimate | Confidence Interval* | Sig. |
|---|---|---|---|
| (Intercept) | 0.69 | 0.48 – 0.90 | **<0.001** |
| success [Hit] | -0.42 | -0.57 – -0.27 | **<0.001** |
| trial | -0.04 | -0.06 – -0.03 | **<0.001** |

$\sigma^2$ = 0.59; $\tau_{00}$ id = 0.38 (n = 119); ICC = 0.39; 1190 observations

*α = 0.05

**Table S26:** Selected untransformed response time mixed model.

| Covariate | Estimate | Confidence Interval* | Sig. |
|---|---|---|---|
| (Intercept) | 2449.01 | 2033.87 – 2864.16 | <0.001 |
| success [Hit] | -564.51 | -884.80 – -244.23 | 0.001 |
| trial | -32.47 | -65.58 – 0.64 | 0.055 |

*α = 0.05

The shorter times of correct answers support that participants were mostly sure about their answers when they replied correctly, and that they did not succeed in the test by answering randomly. Regarding the effect of trial, it must be attributable to the better comprehension of the experiment that takes place along its development. Note that no interaction between trial and success was found and, thus, that effect of trial in response time was independent from succeeding on that particular trial and vice versa. These findings relate to the absence of effect of trial on success in the simple recognition memory test, both meaning that even trials (and better comprehension of the experiment) elicit shorter response times, trial (and better comprehension of the game) has no significant effect on success.
